# Non-image forming vision as measured through ipRGC-mediated pupil constriction is not modulated by covert visual attention

**DOI:** 10.1101/2023.06.27.546729

**Authors:** Ana Vilotijević, Sebastiaan Mathôt

## Abstract

In brightness the pupil constricts, while in darkness the pupil dilates; this is known as the pupillary light response (PLR). The PLR is driven by all photoreceptors: rods and cones, which contribute to image-forming vision, as well as intrinsically photosensitive retinal ganglion cells (ipRGCs), which contribute to non-image-forming vision. Rods and cones cause immediate pupil constriction upon light exposure, whereas ipRGCs cause sustained constriction for as long as light exposure continues. Recent studies have shown that the initial PLR is modulated by covert attention; however, it remains unclear whether the same holds for the sustained PLR. Here, we investigated the effect of covert attention on sustained, ipRGC-mediated pupil constriction. We leveraged the fact that ipRGCs are predominantly responsive to blue light, causing the most prominent sustained constriction in response to blue light. Replicating previous studies, we found that the pupil constricted more when either directly looking at, or covertly attending to, bright as compared to dim stimuli (with the same color). We also found that the pupil constricted more when directly looking at blue as compared to red stimuli (with the same luminosity); crucially, however, we did *not* find any difference in pupil size when covertly attending to blue as compared to red stimuli. This suggests that ipRGC-mediated pupil constriction, and possibly non-image-forming vision more generally, is not modulated by covert attention.

**Significance statement:** When we think of vision, we generally think of image-forming vision, that is, seeing things. However, vision can also be “non-image-forming”; for example, our day-night rhythm and pupil size are regulated by visual input, but not in a way that gives rise to conscious visual awareness. While visual attention shapes image-forming vision, its influence on non-image forming vision remains unclear. We investigated this by using ipRGCs,which contribute to non-image-forming vision and are responsive to blue light. Aside from replicating the effect of covert attention on image-forming vision, we showed that pupil constriction differed between directly looking at blue/ red stimuli, but not during covert attention to these stimuli. This suggests that non-image forming vision is not influenced by covert visual attention.

## Introduction

The pupil light response (PLR) refers to the constriction of the pupil when exposed to brightness, and the dilation of the pupil when exposed to darkness. The PLR is driven by all photoreceptors: rods, cones, and intrinsically photosensitive retinal ganglion cells (ipRGCs) (Barrionuevo et al., 2023; Mathôt et al., 2014; McDougal & Gamlin, 2015). Rods and cones play a pivotal role in image-forming vision by contributing to the perception of visual details, colors, and shapes (Schmidt & Kofuji, 2008). In contrast, ipRGCs play a crucial role in non-image forming vision, that is, visual functions that occur without conscious visual awareness (Berson, 2003; Mathôt, 2018; Provencio et al., 2000; Schmidt & Kofuji, 2008; Zele et al., 2011).

The contribution of rods and cones, and thus of the image-forming pathway, to the PLR is mainly to the initial phase of pupil constriction that is triggered about 200 ms after the onset of a stimulus. However, the input from rods and cones lasts only for around 1.5-2 s, after which the photoreceptors adapt quickly. The contribution of ipRGCs, and thus of the non-image forming pathway, to the PLR is mainly to the sustained phase of pupil constriction that emerges about five to ten seconds after the onset of a stimulus, around the time that rods and cones start to be heavily adapted. ipRGCs receive input from both rods and cones, but they also possess their own photopigment, called melanopsin, which allows them to respond even without input from rods and cones (Barrionuevo et al., 2023; Berson et al., 2002; Dacey et al., 2005; Gamlin et al., 2007; Gnyawali et al., 2022; Markwell et al., 2010; Mathôt, 2018). ipRGCs are mostly sensitive to blue light because the peak sensitivity of the photopigment melanopsin is around 482 nm (Adhikari et al., 2016; Berson, 2003; Mathôt, 2018). Thus far, ipRGCs have been found to play an important role in regulation of the circadian rhythm: when ipRGCs are exposed to light, particularly in the blue spectrum, they send signals to the suprachiasmatic nucleus (SCN) in the brain, which is considered the central pacemaker of the circadian system (Zele et al., 2011). Importantly, ipRGCs are much less prone to adaptation than rods and cones are (Wong, 2012). In sum, although consistent pupil constriction to a long-lasting stimulus, such as daylight, seems to be a unified response, it actually consists of two components: an initial phase, driven by (image-forming) rods and cones, and a sustained phase, driven by (non-image forming) ipRGCs.

Previous studies have shown that the PLR is susceptible to covert attention (Binda et al., 2013; Mathôt et al., 2013; Naber et al., 2013; Unsworth & Robison, 2017). In a study by Mathôt et al. (2013), participants were presented with a display that was vertically divided into a bright and a dark half. At the same time, participants were cued towards either the bright or the dark side, which predicted the location of an upcoming target. The results showed that the PLR is modulated by covert visual attention: the pupil constricted when covertly attending to the bright side as compared to the dark side, even when visual input and gaze position was constant.

Importantly, this study (as well as other studies on the same research question), focused on measuring only the first two seconds after the cue presentation. This time window captures only the initial phase of the PLR, which is driven by rods and cones. Therefore, this study and others leave open the question of whether the sustained phase of the PLR, which is driven by ipRGCs (and thus by non-image-forming vision), is also susceptible to covert attention.

We addressed this open question in the present study. To do so, we leveraged the fact that ipRGCs are predominantly responsive to blue light, causing the most prominent sustained constriction in response to the blue light. We first conducted a Validation experiment that involved directly looking at isoluminant red or blue displays for a duration of 15 seconds. We observed a distinct decrease in pupil size when participants were directly looking at the blue display, as compared to the red display (Figure 2, upper row, right column). We then proceeded to examine whether this effect is susceptible to covert attention in the Main experiment. We designed a task in which participants covertly attended to a stream of letters. Crucially, we manipulated the brightness (dim/bright) and the color (red/blue) of the placeholders on which the letters were superimposed while measuring participants’ pupil sizes (Figure 1c). We carefully matched the luminance of red and blue stimuli, because any pupil size difference should not result from differences in luminance, which would affect the initial pupil response, but rather from differential activation of ipRGCs. If pupil size is smaller when attending to bright stimuli as compared to dark stimuli (given a constant color), then this would show that rod-and-cone mediated pupil constriction, and likely image-forming vision more generally, is modulated by covert visual attention. If pupil size is smaller when attending to blue stimuli as compared to red stimuli (given a constant brightness), then this would show that ipRGC-mediated pupil constriction, and likely non-image-forming vision more generally, is modulated by visual attention.

**Figure 1.**
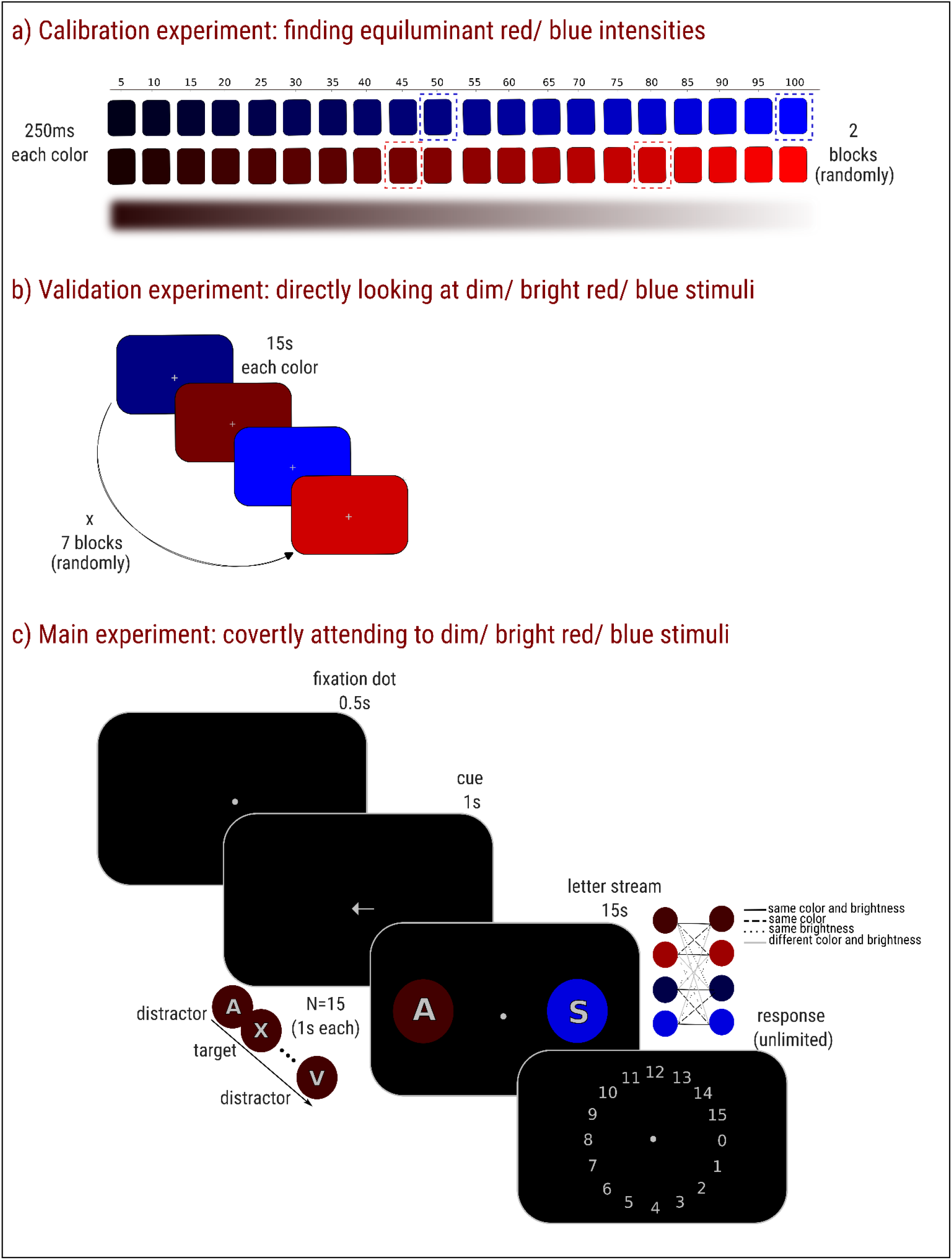
The experimental procedure. *a)* The Calibration experiment. Participants passively looked at the center of the screen while different intensities of red and blue colors were presented in the background. Each color intensity, ranging from 0 to 100 in steps of 5, was presented twice in a random order. Each color intensity was shown for 250 ms followed by a black screen (3000 ms). The outcome of the calibration experiment were participant-specific isoluminant intensities of dim/ bright red and blue. The average color intensities are highlighted by dashed rectangles for illustrative purposes. *b)* The Validation experiment. The color intensities that were obtained in the Calibration experiment were now used in the Validation experiment (and later in the Main experiment). Here, only those four color intensities were presented for 15s. Each color intensity was presented 7 times in a random order. *c)* The Main experiment. Each trial began with a presentation of a fixation dot for .5 s, that was followed by a cue presentation lasting for 1 s. The cue indicated the side to which participants should covertly attend. Finally, two streams of letters appeared on both sides of the screen. Both streams consisted of 15 letters. Only the to-be-attended stream contained the targets (‘X’ letters). The letters on each side were superimposed on placeholder filled circles that were either dim/ bright red/ blue. Participants covertly attended to the indicated stream (to-be-attended stream) and reported the number of targets.

**Figure 2.**
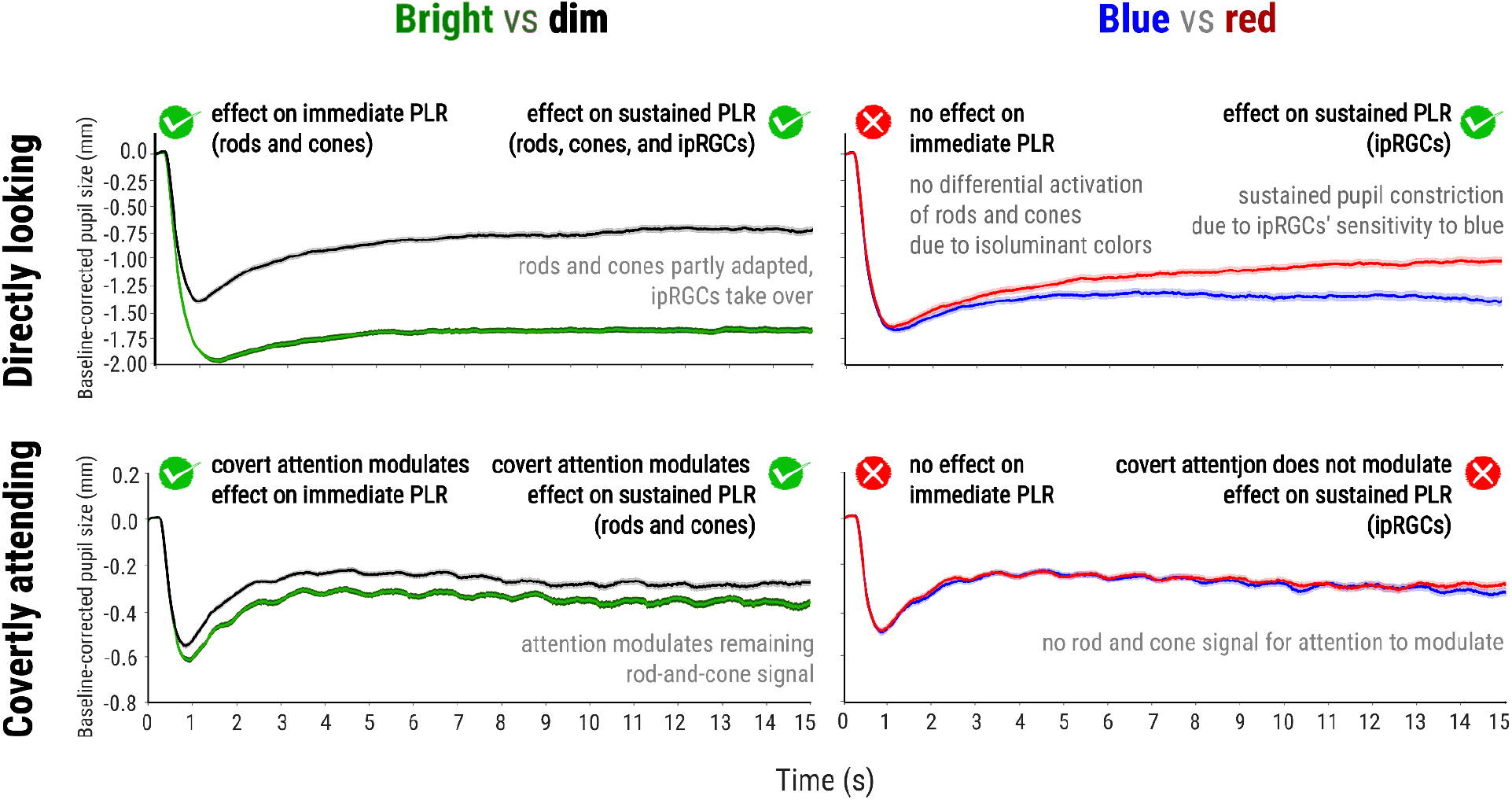
The pupillary results during direct gaze (the Validation experiment; upper row) and covert attention (the Main experiment; lower row). Pupil size when directly looking and covertly attending to bright vs. dim (left column) and blue vs. red (right column). *Note:* Error bars represent the standard error.

## Materials and Methods

All experimental methods and analyses were preregistered here.

### Participants

In total 91 participants, psychology students from the University of Groningen, were recruited and participated in the Calibration and Validation experiments. From these, 30 participants (predetermined sample size) were eligible for participation in the Main experiment upon signing the informed consent form. All participants had normal or corrected-to-normal vision. Participants received course credits for their participation. On the basis of a checklist developed by the Ethical committee (EC-BSS) at the University of Groningen, the study was exempt from full ethical review (PSY-2223-S-0165).

### Apparatus and data acquisition

The experiments were programmed in OpenSesame (Mathôt et al., 2012) using PyGaze for eye tracking (Dalmaijer et al., 2014). Stimuli were presented on a 27-inch monitor (1,920 × 1,080 pixels resolution; refresh rate: 60 Hz) and an EyeLink 1000 (sampling frequency of 1000 Hz; SR Research), was used for eye tracking. Participants’ right eyes were recorded. All experiments were conducted in a dimly lit room.

### Experimental design and procedure

The experimental procedure consisted of three parts: the Calibration, the Validation, and the Main experiment. The first two parts lasted for about 20 minutes and served as an entry point to the Main experiment (see below). An essential aspect of the Main experiment involved orthogonally manipulating the brightness (dim/ bright) and the color (red/ blue) of letters’ placeholders to which participants covertly attended. To ensure that different colors of the same brightness (dim red - dim blue or bright red - bright blue) did in fact not differ in brightness, the Calibration and Validation experiments were participant-specific. That is, the Calibration experiment aimed to determine participant-specific isoluminant intensities of dim/ bright red and blue that were subsequently validated in the Validation experiment and later (for eligible participants) used in the Main experiment. By doing so, we ruled out the possibility that any pupil-size differences observed when presenting colors within the same brightness category were due to variations in luminance between red and blue but rather were due to differential activation of intrinsically photosensitive retinal ganglion cells (ipRGCs).

The Calibration and Validation experiments were conducted on the same day, whereas the Main experiment occurred within a range of 1 to 7 days after. Prior to the start of each part, after making sure that the participant was well seated at about 60 cm distance from the computer, an eye-tracking calibration-validation procedure was run. A chin-rest was used to keep the participant’s head in a stable position. In addition to the eye-tracking calibration-validation procedure, a 1-point eye-tracker recalibration (“drift-correction”) was performed before each trial.

### Calibration experiment: finding isoluminant red/ blue intensities

The goal of the Calibration experiment was to find isoluminant intensities of red and blue. Specifically, we used an automated procedure to determine participant-specific intensities of red and blue that induced the same initial pupil constriction, based on the assumption that this initial constriction is driven only by rods and cones and not by ipRGCs (for similar calibration procedures see Kinzuka et al., 2022; Mathôt et al., 2023; Wardhani et al., 2022). During the Calibration experiment, participants looked passively at a central fixation cross while they were exposed to dim/ bright red/ blue displays that were presented for 250 ms followed by 3000 ms of a black screen (Figure 1a). The stimulus’ intensities varied from 0 to 100% (of the monitor’s maximum brightness) in steps of 5%. Each color intensity was presented twice, and the colors were presented in a random order.

For each display presentation, we determined the strength of pupil constriction by fitting a pupil-constriction template (based on the pupil constriction of the last author) to the pupil response. The fitting procedure had four parameters: a horizontal shift, reflecting the latency of the response; a vertical shift, reflecting baseline pupil size; a horizontal scaling, reflecting the maximum constriction velocity; and a vertical scaling, reflecting the constriction strength. For our purpose, constriction strength was the relevant parameter.

Next, we fitted a second-order polynomial to predict constriction strength from stimulus intensity, separately for red and blue displays. We then determined a pair of red and blue intensities that resulted in the strongest constriction that could be matched, such as 100% blue and 83% red, or 89% blue and 100% red (one of the colors was always 100%). These intensities were used as the bright intensities. Next, we used the 50% blue intensity and the red intensity that matched in terms of constriction strength as the dim intensities, such as 50% and 45% red (blue was always 50%). The final outcome of the Calibration experiment were two (bright, dim) isoluminant intensities for both red and blue.

Participants were eligible for participation in the Main experiment only if the Calibration experiment revealed a systematic and biologically plausible pattern of pupil constriction, based on visual inspection of the data.

### Validation experiment: directly looking at dim/ bright red/ blue stimuli

Once these intensities were obtained, the Validation experiment started, in which participants were exposed to a prolonged presentation (15s) of only the obtained participant-specific intensities of each color, followed by 2500 ms of the black screen, while participants passively fixated the center of the screen (Figure 1b). Each color was presented 7 times, resulting in 28 trials (4 × 7) in total. The validation results served as another entry criterion to the Main experiment: only participants who, based on visual inspection of the data, showed a systematic and biologically plausible pattern of bigger constriction during the presentation of the blue color during direct looking were invited to participate in the Main experiment. Since the decision was made on a level of a single participant (N = 1), no statistical tests were performed and this was decided based on visual inspection.

As described under Participants, only a minority of participants (30 of 91) was invited to participate in the main experiment. This is because perfect calibration is difficult to achieve (e.g. due to blinking or recording artifacts) and yet the quality of the calibration is crucial for the validity of the Main experiment; we therefore chose to include only participants for whom we were sure that the quality of the calibration was very high.

### Main experiment: covertly attending to dim/ bright red/ blue stimuli

Each trial began with a presentation of a fixation dot (0.29°) for .5 s, followed by a cue (1.16°) for 1 s. The cue was an arrow pointing either left or right, and indicating the side to which participants should covertly attend. The cue was 100% valid. Next, two streams of letters (gray letters with a black outline; 2.31°) appeared on both sides of the screen (one-side eccentricity: 13.87° in total). The letters on each side were superimposed on placeholders, which were filled circles (2.89°) that were either dim/ bright red/ blue (Figure 1c). The exact intensities of color-brightness combinations were derived from the Calibration experiment and were participant-specific (the average intensities are highlighted in Figure 1a for illustrative purposes). Both streams (to-be-attended stream and to-be-ignored stream) displayed 15 letters in total, with a pace of 1 s per letter. The streams were not the same, and only the to-be-attended stream contained targets (the letter ‘X’). The color and brightness of the placeholder circles of the to-be-attended and to-be-ignored stream were fully crossed, featuring 16 combinations [4 (*dim red*/ *dim blue*/ *bright red*/ *bright blue*) x 4] in total. The side of the to-be-attended stream was randomly varied. Participants’ task was to covertly attend to the cued stream, count the number of targets that appeared, and report this number at the end of each trial. Each letter had a 20% probability of being a target, resulting in an unpredictable number of targets and a constant hazard rate to encourage sustained attention (i.e. the chance of the next letter being a target was always the same). The experiment consisted of 6 practice (not analyzed) trials and 224 experimental trials. Experimental trials were shuffled and divided in 7 blocks.

### Data preprocessing

Following the workflow for preprocessing pupillary data that we described elsewhere (Mathôt & Vilotijević, 2022), we first interpolated blinks and downsampled the data by a factor of 10. Next, we converted pupil size measurements from arbitrary units to millimeters of diameter by using the formula specific to our lab (Wilschut & Mathôt, 2022). Finally, we baseline-corrected the data by subtracting the mean pupil size during the first 50 ms after the onset of the Letter stream (baseline period) from all subsequent pupil-size measurements on a trial-by-trial basis.

### Data exclusion

First, we checked whether participants made eye movements and excluded trials in which the deviation from the center of the screen was larger than 6.93° (halfway to the letter stream) and lasted longer than 10 ms (681 trials excluded). Trials containing baseline pupil sizes of ±2 z-scores were considered outliers, and hence excluded from the data (442 trials excluded). In total, 1123 trials (16.71 %) were excluded from the data.

## Results

### Validation: directly looking at dim/ bright red/ blue stimuli

First, we used the results from the Validation experiment to analyze pupil size changes when participants were directly looking at dim/bright red/blue. We ran two Linear Mixed-Effects (LME) analyses to investigate the effect of brightness (dim vs. bright) and, separately, the effect of color (red vs. blue) on the pupil. We used the entire 15s trace as a window of interest. For the analysis that looked into the effect of brightness on the pupil, we aggregated data across colors, so that both dim red and dim blue trials were coded as ‘dim’ and contrasted to both bright red and bright blue trials, which were coded as ‘bright’. Our model included Mean pupil size as a dependent variable, Brightness (dim vs. bright) as a fixed effect and by-participant random intercepts and slopes. Similarly, for the analysis that looked into the effect of color on the pupil, we aggregated data across brightness, so that both dim red and bright red trials were coded as ‘red’ and contrasted to both dim blue and bright blue trials, which were coded as ‘blue’. Our model included Mean pupil size as dependent variable, Color (red vs. blue) as a fixed effect and by-participant random intercepts and slopes.

We found a main effect of brightness (*b* = .84, *SE* = .03, *t* = 29.27, *p* < .001), reflecting that the pupil constricted more when directly looking at bright as compared to the dim displays. We also found a main effect of color (*b* = .19, *SE* = .04, *t* = 4.63, *p* < .001), reflecting that the pupil constricted more when directly looking at the blue trials as compared to the red trials (Figure 2 upper row).

Qualitatively, we also observed the typical pattern of an initial equivalent constriction to blue and red, which reflects that we had successfully matched the intensities of red and blue, followed by a gradually emerging difference such that constriction is more sustained when looking at a blue as compared to a red display. This pattern is the behavioral hallmark of the involvement of ipRGCs (Mathôt, 2018; Barrionuevo et al., 2023).

### Main experiment: covertly attending to dim/ bright red/ blue stimuli

Next, we used the results from the Main experiment to analyze pupil size changes when participants were covertly attending to dim/ bright red/ blue. To test this, we used the python library time_series_test, which combines cross-validation with linear mixed effects (LME) modeling in a way that is suitable for pupil-size data (Mathôt & Vilotjevi ć, 2022)^1^. The idea behind it is that the cross-validation localizes the sample at which the effect is strongest within the full letter stream time window, and then conducts a single LME analysis on these samples. We ran this analysis two times to investigate the effect of brightness (bright vs. dim) and the effect of color (red vs. blue) on the pupil. For the analysis that looked into the effect of brightness on the pupil, we selected only the trials in which the to-be-attended and to-be-ignored streams’ placeholder had the same color (same color trials; see Figure 1c). Next, we aggregated data across colors, so that both dim red and dim blue trials were coded as ‘dim’ and contrasted to both bright red and bright blue trials, which were coded as ‘bright’. Our model included Mean pupil size as a dependent variable, Brightness (dim vs. bright) as a fixed effect and by-participant random intercepts and slopes. We used the entire 15s trace as a window of interest. Similarly, for the analysis that looked into the effect of color on the pupil, we selected only the trials in which to-be-attended and to-be-ignored streams’ placeholder had the same level of brightness (same brightness trials; see Figure 1c). Next, we aggregated data across brightness, so that both dim red and bright red trials were coded as ‘red’ and contrasted to both dim blue and bright blue trials, which were coded as ‘blue’. Our model included Mean pupil size as dependent variable, Color (red vs. blue) as a fixed effect and by-participant random intercepts and slopes.

We found a main effect of brightness (*z* = 5.49, *p* < .001, tested at samples 170, 460 and 190), suggesting that the pupil constricted more when covertly attending to bright as compared to dim placeholders. However, we did not find a main effect of color, (*z* = 0.73, *p* = .462, tested at samples 1560, 1460 and 190), suggesting that there were no differences in pupil constriction when covertly attending to blue as compared to red placeholders (Figure 2 lower row) (for more detailed differences see Appendix).

To rule out the effect of task difficulty on pupil size, we checked the accuracy across different conditions. We ran a Generalized Linear Mixed-Effects (GLME) analysis, including Accuracy as a dependent variable, Color (blue vs. red), Brightness (bright vs. dim) of the covertly attended stream, and their interactions as a fixed effects and by-participant random intercepts and slopes for each fixed effect. We found no main effects (Color: *b* = -.25, *SE* = .16, *t* = -1.54, *p* = .123; Brightness: *b* = -.15, *SE* = .15, *t* = -.97, *p* = .329), nor the interaction (*b* = .02, *SE* = .16, *t* = .18, *p* = .135), suggesting that the accuracy did not differ across different conditions (Figure 3).

**Figure 3.**
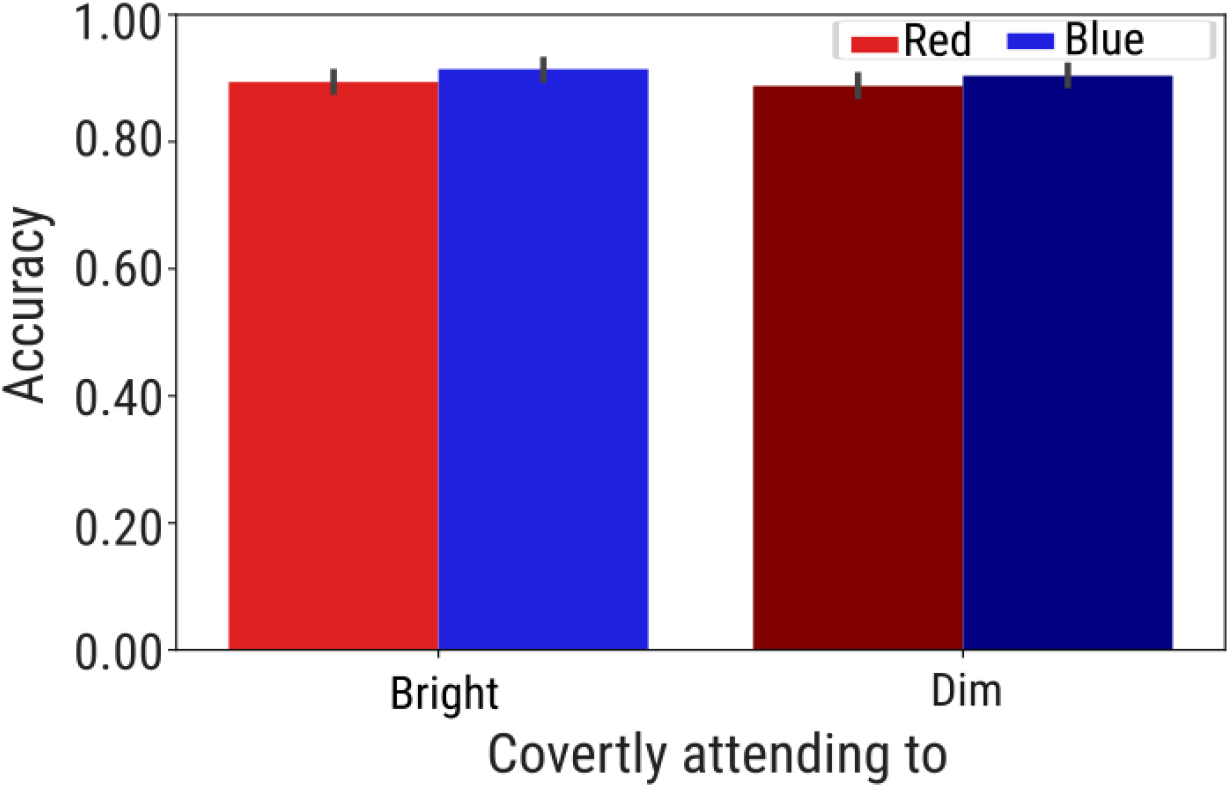
The behavioral results. Accuracy as a function of color and brightness of the covertly attended stream. *Note:* Error bars represent the standard error.

**Figure 4.**
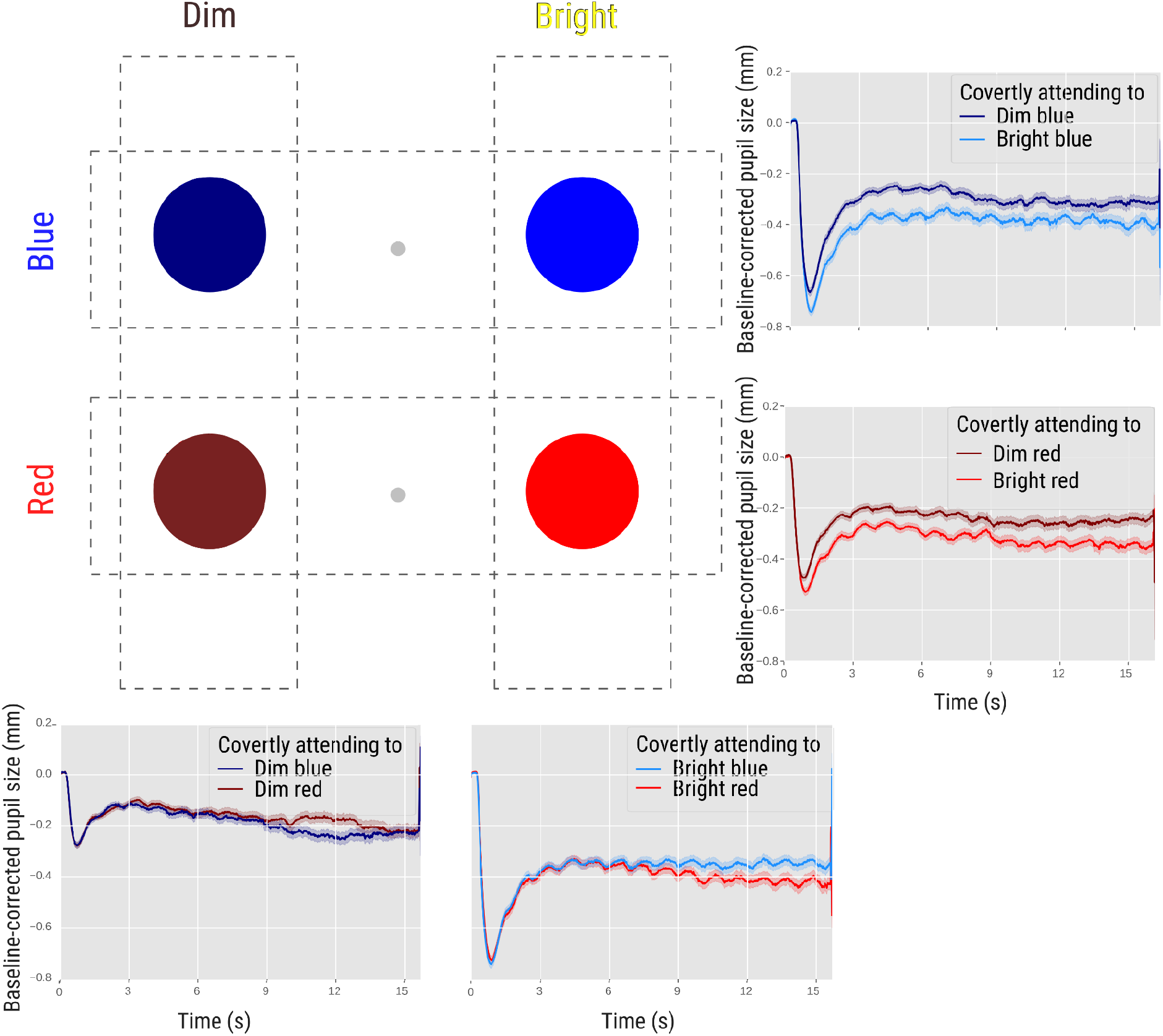
Separate comparisons. Pupil size when covertly attending to bright/ dim blue/ red under the same color (rows)/ brightness (columns) conditions.

## Discussion

In our study, we investigated the effect of covert attention on the sustained phase of the pupillary light response (PLR), which is mediated by intrinsically photosensitive retinal ganglion cells (ipRGCs); we compared this to the well-established effect of covert attention on the initial PLR (Binda et al., 2013; Mathôt et al., 2013; Naber et al., 2013), which is mediated by rods and cones. To do so, we exploited the fact that ipRGCs are most responsive to blue light, resulting in a prominent and sustained constriction when exposed to blue light stimuli. We found that the pupil constricted more when either directly looking at or covertly attending to bright as compared to dim stimuli (with the same color), replicating the effect of covert attention on the initial phase of the PLR. We also found that the pupil constricted more when directly looking at blue as compared to red stimuli (with the same luminosity); crucially, however, we did *not* find any difference in pupil size when covertly attending to blue as compared to red stimuli.

The effect of brightness (bright vs. dim) on the initial phase of the PLR, both when looking directly at a stimulus and when covertly attending to it, is illustrated in the left column of Figure 2. Crucially, there is no effect of color on the initial phase of the PLR, neither when looking directly at a stimulus nor when covertly attending to it, as illustrated in the right column of Figure 2; this is due to the fact that the we (successfully) controlled the influence of rods and cones by equating the luminance of the colors in order to isolate ipRGC activation during the sustained pupil response. During the direct looking condition, there was an effect of color (blue vs. red) on the sustained phase of the PLR; the activation of ipRGCs is slow, and as a result, this effect emerged only after approximately five seconds. As time progresses and ipRGCs become increasingly active, this difference becomes more pronounced, reaching approximately 0.4 mm towards the end (Figure 2 right column, upper row).

The crucial finding of the present study is shown Figure 2 right column, lower row. This shows that covertly attending to (unlike directly looking at) blue as compared to red does *not* result in increased sustained pupil constriction. This suggests that the sustained phase of the PLR, which is driven by ipRGCs, is impervious to covert attention.

The fact that we observed a persistent pupil size difference when covertly attending to bright/ dim stimuli may seem to conflict with the general idea, which we have also reiterated here, that rods and cones only contribute to the initial phase of the PLR, and that sustained pupil constriction is driven ipRGCs (Lucas et al., 2020; Mathôt, 2018; Park et al., 2011; Thompson et al., 2005). If this is correct, then should we not observe that the effect of covertly attending to bright/ dim stimuli is short-lived and disappears after several seconds as rods and cones adapt and ipRGCs take over? A likely explanation for the sustained effect that we observed here is that rods and cones did not undergo complete adaptation. Specifically, participants were continuously exposed to a stream of letters overlaid on static placeholders. With the presentation of each new letter, participants’ eyes naturally exhibited microsaccades. Both the letter changes and these microsaccades resulted in refreshing of rod and cone activation, especially because our placeholders had sharp edges that are continuously carried into and out of receptive fields. In other words, the sustained phase of the PLR could still be driven by rods and cones, which were repeatedly activated. Future research could test the effect of covert attention on ipRGCs while ensuring full adaptation of rods and cones. This could be accomplished using large and static stimuli without sharp edges, which would prevent microsaccades from re-activating rods and cones.

In conclusion, we replicated the effect of covert attention on the initial phase of the PLR, which is driven by rods and cones. However, we did not find any difference in pupil size when covertly attending to blue as compared to red stimuli (with the same luminosity), whereas we did observe this difference when participants directly looked at the same blue or red stimuli. This suggests that the sustained phase of the PLR, which is driven by ipRGCs, is not modulated by covert attention. This finding has important implications for our understanding of non-image-forming vision, that is, for visual functions that are not accompanied by visual awareness, such as regulation of circadian rhythm (Schmidt & Kofuji, 2008; Zele et al., 2011). Since ipRGCs are strongly involved in non-image-forming vision (Güler et al., 2008; Mathôt, 2018; Schmidt & Kofuji, 2008), our results imply that the pathway that underlies non-image-forming vision may—unlike image-forming vision—not be modulated by covert visual attention.

### Open-practices statement

Experimental data, the experiment, and analysis scripts can be found here.

## Acknowledgments

This research was supported by the Innovational Research Incentives Scheme VIDI (VI.Vidi.191.045) from the Dutch Research Council (NWO) to Sebastiaan Mathôt.

## Appendix

## Footnotes

1 Since this analysis is very time-consuming, we downsampled the data once again by a factor of 10.

